# Pontine waves accompanied by short hippocampal sharp wave-ripples during non-rapid eye movement sleep

**DOI:** 10.1101/2022.06.03.494781

**Authors:** Tomomi Tsunematsu, Sumire Matsumoto, Mirna Merkler, Shuzo Sakata

## Abstract

Ponto-geniculo-occipital (PGO) or pontine (P) waves have long been recognized as an electrophysiological signature of rapid eye movement (REM) sleep. However, P-waves can be observed not just during REM sleep, but also during non-REM (NREM) sleep. Recent studies have uncovered that P-waves are functionally coupled with hippocampal sharp wave-ripples (SWRs) during NREM sleep. However, it remains unclear to what extent P-waves during NREM sleep share their characteristics with P-waves during REM sleep and how the functional coupling to P-waves modulates SWRs. Here, we address these issues by performing multiple types of electrophysiological recordings and fiber photometry in both sexes of mice. P-waves during NREM sleep share their waveform shapes and local neural ensemble dynamics at a short (∼100 ms) timescale with their REM sleep counterparts. However, the dynamics of mesopontine cholinergic neurons are distinct at a longer (∼10 s) timescale: although P-waves are accompanied by cholinergic transients, the cholinergic tone gradually reduces before P-wave genesis during NREM sleep. While P-waves are coupled to hippocampal theta rhythms during REM sleep, P-waves during NREM sleep are accompanied by a rapid reduction in hippocampal ripple power. SWRs coupled with P-waves are short-lived and hippocampal neural firing is also reduced after P-waves. These results demonstrate that P-waves are part of coordinated sleep-related activity by functionally coupling with hippocampal ensembles in a state-dependent manner.

## Introduction

Sleep is a fundamental homeostatic process in the animal kingdom. Determining its characteristics and exploring its functions is essential in neuroscience. Sleep states in mammals can be determined by multiple electrophysiological markers. For example, in addition to low muscle tone, cortical electroencephalograms (EEGs) during NREM sleep are characterized by slow and delta (<4 Hz) oscillations. Additionally, other oscillations including sleep spindles and hippocampal sharp wave-ripples (SWRs) have been extensively studied ^1–6^.

While theta (∼7 Hz) oscillations are prominent during REM sleep, ponto-geniculo-occipital (PGO) waves have long been recognized ^7–11^: PGO waves during REM sleep were initially reported in cats^12^. Subsequently, similar sub-second brain waves have been observed in other mammalian species, including rats ^8, 13^, mice ^14, 15^, macaques ^16, 17^ and humans ^18^. In rats, PGO waves are often called pontine-waves or “P-waves” ^19^.

P-waves consist of ∼100 ms waves originating from mesopontine cholinergic nuclei, such as the pontine tegmental nucleus, laterodorsal tegmental nucleus and sublaterodorsal nucleus in rodents ^8, 14, 15, 20^, which play an imperative role in REM sleep induction ^21–25^. P-waves can be observed across a wide range of brain regions, not just the visual pathway, but also in limbic areas ^26^ and the cerebellum ^27, 28^.

During REM sleep, P-waves and hippocampal theta oscillations are coupled ^15, 16, 29, 30^: the timing of P-waves is locked to a certain phase of ongoing theta oscillations. While this was originally discovered in cats ^29, 30^, recent studies confirmed a similar phase-coupling in mice ^15^ and macaques^16^. P-waves can also be observed during NREM sleep although the frequency of P-waves is less ^15, 16^. Intriguingly, recent two studies reported that P-waves during NREM sleep are coupled with hippocampal SWRs, implying a role in systems memory consolidation ^15, 16^.

Although these results suggest brain-wide coordination of sleep-related neural events across sleep states, the following issues remain unsolved. First, how similar and different are P-waves during NREM sleep compared to those during REM sleep? While the detection of P-waves relies on local field potentials, the underlying neural dynamics may be different. However, no previous studies have addressed this issue directly. Second, it remains to be determined what is the consequence of the functional coupling between SWRs and P-waves with respect to hippocampal ensembles. Because the duration of SWRs is a critical factor for memory consolidation ^31^, it is important to determine how the duration of SWRs and the excitability of hippocampal neurons are influenced by P-waves. In the present study, by analyzing our published datasets ^14, 15^ and performing additional electrophysiological and fiber photometry experiments in mice, we report that although brainstem neural activity associated with P-waves is similar between two sleep states, mesopontine cholinergic activity exhibits distinct dynamics at a long (∼10 s) timescale. We also uncover a non-trivial antagonistic role of P-waves with SWRs, suggesting the functional importance of P-waves in systems memory consolidation.

## Methods

### Animals

24 mice (8-37 weeks old, 19 males, 5 females) were used in this study and were housed individually with a 12 h:12 h light/dark cycle (light on hours: 7:00-19:00 for 22 mice, 8:00-20:00 for 2 mice). Mice had *ad libitum* access to food and water. Their genotypes consisted of wild-type, ChAT-IRES-Cre (JAX006410), or ChAT-IRES-Cre::Ai32 (JAX012569) on a C57BL/6 background. For brainstem silicon probe recordings, 7 animals were used (9 recordings). For hippocampal silicon probe recordings, 2 animals were used (4 recordings). For hippocampal Neuropixels probe recordings, 3 animals were used (3 recordings). For P-wave recordings with bipolar electrodes, 8 animals were used (15 recording sessions with 9 bilateral recordings). For photometry recordings, 4 animals were used (8 recordings). All experiments were performed during the light period (zeitgeber time (ZT) 1-10). All experimental procedures were performed in accordance with the United Kingdom Animals (Scientific Procedures) Act of 1986 Home Office regulations and approved by the Home Office (PPL 70/8883) and the Animal Care and Use Committee of Tohoku University (approval no. 2019LsA-018). While the majority of the datasets were taken from previous studies ^14, 15^, the following datasets were added for the present study: 4 recordings (2 mice) for bilateral P-wave recordings; 3 recordings (3 mice) for Neuropixels probe recordings; 2 recordings (2 mice) for photometry recordings.

### Surgical procedures

Detailed procedures were described elsewhere ^15^. Briefly, mice were anesthetized with isoflurane (2-5% for induction, 1-2% for maintenance) and placed in a stereotaxic apparatus (SR-5M-HT or SR-6M-HT, Narishige). Body temperature was maintained at 37°C with a feedback temperature controller (40–90–8C, FHC). After exposing the skull, two bone screws were implanted on the skull for cortical EEGs in frontal cortical regions (n = 18 mice) or parietal cortical regions (n = 2 mice). Twisted wires (AS633, Cooner Wire) were inserted into the neck muscle for EMG. For pontine EEG recording, bipolar electrodes for pontine EEGs were typically implanted in the medial parabrachial nucleus of the pons (5.1 mm posterior, 1.2 mm lateral from bregma, 3.2 mm depth from brain surface) or the sublaterodorsal nucleus (5.1 mm posterior, 0.6 mm lateral from bregma, 2.6 mm depth from brain surface). The bipolar electrodes consisted of 75 or 100 µm diameter stainless wires (FE631309, Advent Research Materials; FE205850, Goodfellow; EKCSS-010, Bio Research Center). The tips of two glued wires were separated by approximately 0.5 mm vertically to differentiate signals. A pair of nuts was also attached on the skull with dental cement as a head-post. After the surgery, Carprofen (Rimadyl, 5 mg/kg) was administered intraperitoneally. The animals were left to recover for at least 5 days. During the habituation period, the animals were placed in a head-fixed apparatus, by securing them by the head-post and placing the animal into an acrylic tube. This procedure was continued for at least 5 days, during which the duration of head-fixed was gradually extended from 10 to 120 min. After this period, a recording was performed for up to 5 hours (see below).

For silicon probe or Neuropixels probe recording, in addition to the procedures above, a day after the habituation period, the animals were anesthetized with isoflurane and a craniotomy to insert a probe into the brainstem or hippocampus was performed. A craniotomy on the left hemisphere (4.0 mm to 5.5 mm posterior, 1.0 to 1.3 mm lateral from bregma) for the brainstem recording or on the left hemisphere (2.0 mm posterior, 1.5 mm lateral from bregma or 2.5 to 3.5 mm posterior, 2.4 to 3.4 mm lateral from bregma) for the hippocampus recording were performed, respectively. To protect and prevent the brain from drying, the surface was covered with biocompatible sealants (Kwik-Sil and Kwik-Cast, WPI). The following day, the animals were placed in the head-fixed apparatus for recording.

For simultaneous in vivo electrophysiology and fiber photometry ^14^, the viral vector (AAV5-CAG-flex-GCaMP6s-WPRE-SV40, Penn Vector Corel titer 8.3 x 10^12^ GC/ml) was injected in the pedunculopontine tegmental nucleus (PPN) and laterodorsal tegmental nucleus (LDT) (-4.5 mm posterior, 1.m mm lateral from bregma, 3.25 mm depth from brain surface). A hybrid implant consisting of a bipolar electrode glued to an optic fiber was implanted 3 mm deep from the brain surface in addition to bone screws for cortical EEGs and twisted wires for EMG. All components were fixed with dental cement.

### in vivo electrophysiological recording

Electrophysiological recordings were performed in a single-walled acoustic chamber (MAC-3, IAC Acoustics) with the interior covered with 3 inches of acoustic absorption foam. The animals were placed in the same head-fixed apparatus, by securing them by the head-post and placing the animal into an acrylic tube. During the entire recording, the animals were not required to do anything actively. For pontine EEG recording, cortical EEG, EMG and pontine EEG signals were amplified (HST/32V-G20 and PBX3, Plexon), filtered (0.1 Hz low cut), digitized at a sampling rate of 1 kHz and recorded using LabVIEW software (National Instruments). A subset of recordings was performed using an Intan Technologies system (RHD 16-channel bipolar-input recording headstage, RHD2000). The recording was performed for approximately 5 hrs between ZT1-6.

For silicon probe recording, a 32-channel 4-shank silicon probe (A4 x 8-5 mm-100-400-177 for brainstem recording or Buzsaki32 for hippocampus recording) was inserted slowly with a manual micromanipulator (SM-25A, Narishige) into the brainstem (3.75 mm – 4.3 mm from the brain surface) or the hippocampus (1.55 mm – 1.85 mm from brain surface). Broadband signals were amplified (HST/32V-G20 and PBX3, Plexon) relative to a screw on the cerebellum, filtered (0.1 Hz low cut), digitized at 20 kHz and recorded using LabVIEW software (National Instruments). For Neuropixels probe recording, the probe was inserted slowly with a motorized manipulator (DMA-1511, Narishige) through the hippocampus (4 mm from the brain surface). Signals were amplified and digitized in the probe and recorded using SpikeGLX (Janelia Research Campus). The recording session was initiated > 1 hr after the probe was inserted into its target depth, to stabilize neural signals. For verification of probe tracks, the rear of the probe was painted with DiI (∼10% in ethanol, D282, Invitrogen) or CM-DiI (0.1% w/v in ethanol, C7001, Invitrogen) before insertion. Pupillometry was also measured simultaneously from a subset of recordings. The experimental procedures and results of pupillometry were reported elsewhere ^15^.

### Simultaneous in vivo electrophysiology and fiber photometry

Detailed procedures were described elsewhere ^14, 15^. Briefly, recordings were performed in an open box (21.5 cm x 47 cm x 20 cm depth) lined with absorbent paper, bedding, and soft food. Electrophysiological signals were recorded at 1 kHz using an interface board (RHD2000, Intan Technologies) and connected to the mouse via an amplifier (RHD2132, Intan Technologies). The fiber photometry system consisted of two excitation channels: a 470 nm LED for Ca^2+^-dependent signals and a 405 nm LED for Ca^2+^-independent isosbestic signals. LED pulses were alternately turned on and off at 40 Hz. The illumination power was adjusted at the tip of the fiber to 0.7-1.37 mW/mm^2^ for the 470 nm LED and 0.4-0.94 mW/mm^2^ for the 405 nm LED. Emission signals were collected back through the optic fiber and directed to a photodetector (NewFocus2151, Newport). A data acquisition module (NI USB-6211, National Instruments) and a custom-written program (LabVIEW) were used to control the LEDs and acquire both fluorescence signals and electrophysiological signals at 1 kHz.

### Histological analysis

After electrophysiological experiments, animals were deeply anesthetized with a mixture of pentobarbital and lidocaine and perfused transcardially with saline or phosphate buffer saline (PBS) followed by 4% paraformaldehyde/0.1 M phosphate buffer, pH 7.4. The brains were removed and immersed in the above fixative solution overnight at 4°C and then immersed in 30% sucrose in PBS for at least 2 days. The brains were cut into coronal or sagittal sections with a sliding microtome (SM2010R, Leica) or with a cryostat (CM3050, Leica) with a thickness of 50 or 100 µm. To identify the location of electrode positions, the brain sections were incubated with NeuroTrace 500/525 Green-Fluorescence (1/350, Invitrogen), NeuroTrace 435/455 Blue-Fluorescence (1/100, Invitrogen) or DAPI (1/1000, Invitrogen) in PBS for 20 min at room temperature (RT). After staining, sections were mounted on gelatin-coated or MAS-coated slides and cover-slipped with 50% glycerol in PBS. The sections were examined with a fluorescence microscope (BZ-9000, Keyence). For Neuropixels probe experiments, the track of the probe was estimated using publicly available software (https://github.com/cortex-lab/allenCCF).

### Data analysis

#### Sleep scoring

Vigilance states were visually scored offline according to standard criteria as described elsewhere ^14, 15^. Wakefulness, NREM sleep, or REM sleep were determined every 4 seconds based on cortical EEG and EMG signals using a custom-made MATLAB GUI (https://github.com/Sakata-Lab/SleepScore) or SleepSign (KISSEI COMTEC). While most cortical EEGs were recorded from frontal cortical regions, the power spectral density was distinct across states (**Supplementary Figure 1**).

#### Photometry signal processing

Detailed procedures were described elsewhere ^14^. Briefly, the median fluorescent signals during each LED pulse were calculated, meaning that the original signals were effectively down-sampled to 40 Hz. Photobleaching was estimated by fitting a single exponential curve and the fluorescent signals were subtracted from the estimate. After applying a low-pass (<4 Hz) filter, the 405 nm signals were scaled based on the 470 nm signals using linear regression. Finally, the 470 nm signals were subtracted from the scaled 405 nm signals.

#### P-wave detection

To detect P-waves, two EEG or LFP signals in the pons were subtracted and filtered (5-30 Hz band-pass filter). If the signals cross a threshold, the event was considered to be a P-wave candidate. To determine the detection threshold, a segment of signals was extracted from the longest NREM sleep episode for a reliable estimation of the noise level. The noise level was estimated by computing root-mean-square (RMS) values in every 10 ms window. The threshold was defined as the mean + 5 x standard deviation of the RMS values. To eliminate potential moving artifacts, EMG signals were also assessed in parallel. If the RMS values of EMG signals cross a threshold, the candidate was discarded. The threshold was defined as the mean + 3 × standard deviation (SD) of the RMS values of EMG signals with 10 ms resolution. The timing of P-waves was defined as the timing of the negative peak.

#### SWR detection

To detect ripples in the hippocampus, LFP signals in CA1 were band-pass filtered at 80-250 Hz with a 3^rd^-order Butterworth filter. The envelope of the filtered signals was calculated by computing root-mean-square values using a sliding window of 20-ms data points. The mean and SD of this “ripple” power profile were used to compute two thresholds (mean + 5×SD and mean + 2×SD), which were used to detect SWRs and to determine their onset, peak timing and offset. To detect SWRs, the period of the ripple power larger than the threshold (mean + 5×SD) was recognized as SWR candidates. The peak timing of each candidate was defined as SWR timing. The onset and offset of each candidate were determined as time points when the candidate’s power profile crossed the threshold (mean + 2×SD). If the onset and offset were overlapped across multiple SWR candidates, they were merged as a single SWR and the SWR timing was defined as the time point providing the maximum power. Finally, candidates shorter than 20 ms or longer than 200 ms were excluded. This process was applied to all silicon probe channels. Signals from the channel which provided the highest frequency of SWRs during NREM sleep, but low frequency of SWRs during REM sleep was used. In Neuropixels probe recording, the position of the CA1 pyramidal cell layer was estimated by computing the ratio between SWR frequency during NREM and REM sleep. The channel that showed the maximum value was chosen.

#### Phase and band power analysis

The phase analysis (**Figure 4B**) was the same as that described previously ^15, 32^. LFP signals from two separate channels were subtracted to minimize volume conduction. To derive band-limited signals at theta band (4 – 10 Hz), a Kaiser finite impulse response filter with sharp transition bandwidth of 1 Hz, pass-band ripple of 0.01 dB and stop-band attenuation of 50 dB. For filtering, MATLAB *filtfilt* function was used. The instantaneous phase of the theta band was estimated from the Hilbert transform and the phase of P-wave occurrence was computed. To quantify the relationship between P-waves and LFP phase, the number of P-waves elicited in each phase bin was calculated. The resultant vector length was computed as an absolute value of the mean resultant vector *r* by using *circ_r* function of CircStats Toolbox ^33^. Rayleigh’s test for non-uniformity of circular data was performed to assess the statistical significance (*p* < 0.05) of the non-uniformity of the P-wave vs LFP phase distribution.

To estimate theta powers (**Figure 4C**), band-limited signals at theta band (4 – 10 Hz) were used. To estimate ripple powers (**Figure 4E**), band-limited signals were derived at 80 – 250 Hz. Then, those signals were cross-correlated with P-wave timing during REM sleep for the theta band and NREM sleep for the ripple band.

Continuous wavelet transform (CWT) (**Figure 4B**) was performed to visualize event-triggered hippocampal LFP signals in the spectro-temporal domain. The CWT was computed using the analytic Morse wavelet with the symmetry parameter, ɣ, and the time-bandwidth product. We used the *cwt* function in MATLAB, which also imlplements L1 normalization.

#### Spectral Inference Criterion analysis

The motivation for this analysis and an algorithm of the spectral inference criterion (SIC) were described in previous studies ^16, 34^ and outlined in **Supplementary Figure 9**. To compute the spectral dependency ratio (SDR) from *X* to *Y*, hippocampal LFPs and pontine LFPs were extracted in a ± 250 ms window from P-wave timing. The spectral power density (PSD) of *X* (hippocampal LFPs) and *Y* (pontine LFPs) was estimated as *SXX* and *SYY* using Welch’s method (*pwelch* function in MATLAB). Then the SDR from *X* to *Y* was calculated as

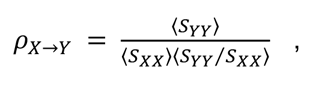

where *ρ_X→Y_* is the SDR from *X* to *Y* and 〈·〉 is the average of the estimated PSD. If *ρ_X→Y_* is larger than *ρ_Y→X_*, then the directional influence from *X* to *Y* is stronger. We computed SDR across all P-wave events in each sleep state.

#### Spike train analysis

For both conventional silicon probe and Neuropixels recording data, Kilosort ^35^ was used for automatic processing, followed by manual curation using phy (https://github.com/cortex-lab/phy). Clusters with ≥20 isolation distance were recognized as single units. The position of recorded neurons was estimated based on histological analysis. This estimation was refined by computing the ratio of SWR frequency between NREM and REM sleep (the channel exhibiting the highest ratio was set as the CA1 pyramidal layer). In the present study, we analyzed only CA1 neurons. All subsequent analyses were performed using custom-written codes (MATLAB, Mathworks).

#### Asymmetry index and modulation index

Asymmetry index (**Figure 4E**) and modulation index (**Figure 4H**) were defined as (*X_post_ − X_pre_)⁄(X_post_ + X_pre_*), where *X_pre_* and *X_post_*are values (ripple power in **Figure 4E**; firing rate in **Figure 4H**) before and after P-wave timing in a certain time window (250 ms), respectively.

#### Statistical analysis

Data were presented as mean ± SEM unless otherwise stated. Statistical analyses were performed with MATLAB. In **Figures 2A and 2B**, Wilcoxon signed-rank test was performed. In **Figure 4F**, Wilcoxon rank-sum test was performed. In **Figure 2D**, linear mixed-effects modeling was performed to analyze the correlation between the response variable (sleep episode duration, *y*) and the predictor variable (P-wave frequency, *X*) while accounting for potential variations across different recordings. The model was specified as *y* = *Xβ* + *Zb* + *ε*, where *y* is the response variable (sleep episode duration), *X* is the fixed effect predictor variable (P-wave frequency), and *β* is a fixed-effects coefficient. *Z* and *b* represent a random effect term for the estimation of group-specific intercepts, accounting for variations across recordings. *ε* represents an error. To estimate the model, MATLAB’s *fitlme* function was used. In **Figures 3B, 4E** and **4H**, Student’s *t*-test was performed.

## Results

### P-waves during NREM and REM sleep

We performed three types of electrophysiological recordings: firstly, we performed pontine EEG recording to detect P-waves by chronically implanting a bipolar electrode (*n* = 11 sessions from 6 animals) (**Figure 1A**). Secondly, we performed silicon probe recording in mid-hindbrain regions (we call it as the “brainstem” afterward) to monitor spiking activity associated with P-waves (*n* = 9 sessions from 7 animals) (**Figure 1D**). Finally, we performed conventional silicon or Neuropixels probe recording in the hippocampal CA1 while monitoring pontine EEGs to investigate the relationship between P-waves and hippocampal oscillations (*n* = 7 sessions from 5 animals) (**Figure 4A**). Across all experiments, we also monitored cortical EEGs and electromyograms (EMGs) to determine sleep states. While we carried out all experiments in a head-fixed condition since we combined silicon probe recording and pupillometry, the sleep architecture was generally comparable with that in a tethered unrestrained condition as reported previously ^15^.

**Figure 1.**
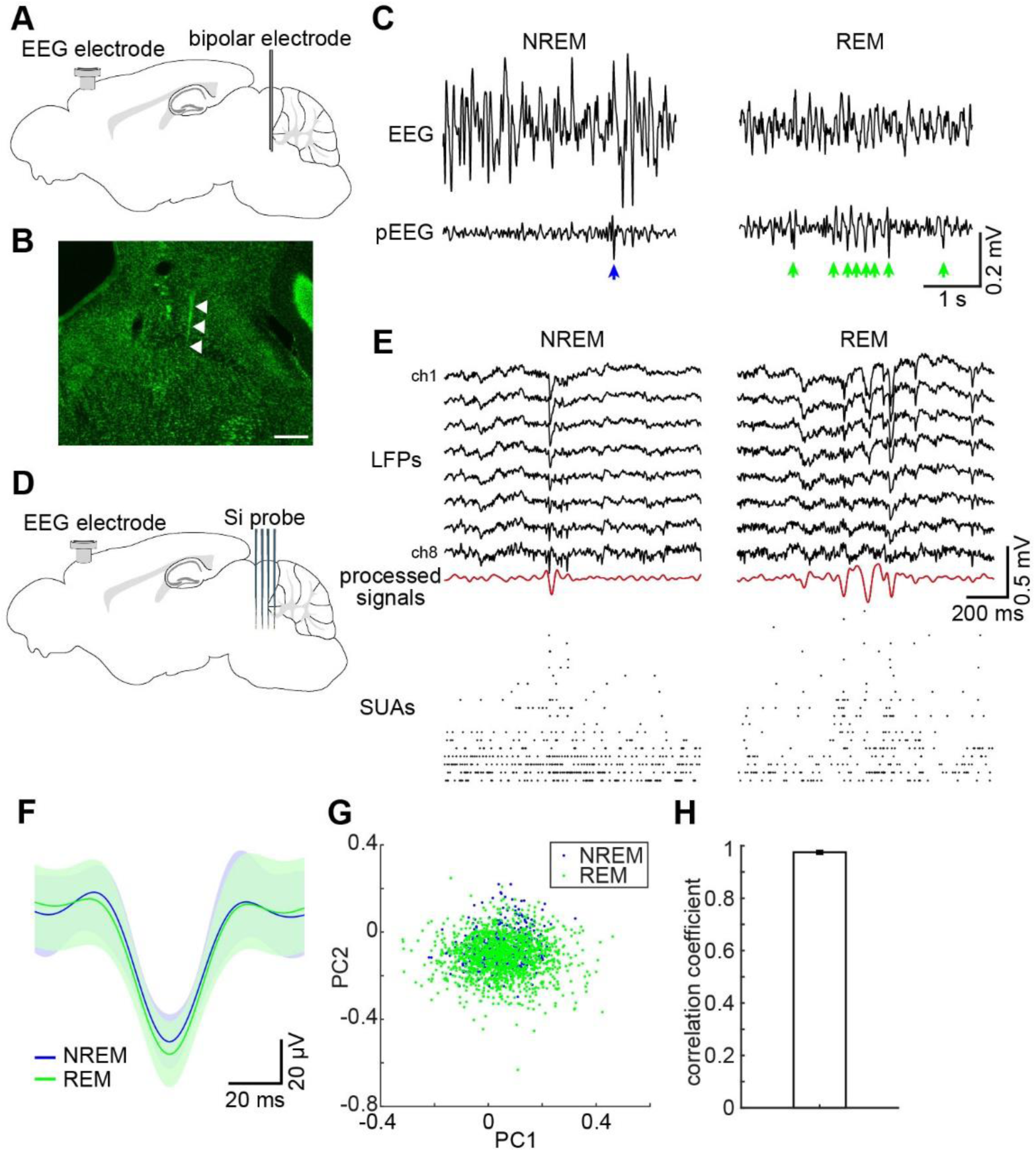
P-waves during NREM and REM sleep. **(A)** A diagram of the experimental approach for P-wave recording, showing a cortical EEG electrode and a bipolar electrode for pontine EEG recording in the mouse. For EMG recording, a twisted bipolar electrode was also inserted in the neck muscle. **(B)** A photograph showing a track of a bipolar electrode in the mesopontine region. Scale bar, 500 µm. **(C)** Examples of P-waves during NREM (*left*) and REM sleep (*right*). **(D)** A diagram of the experimental approach to brainstem population recording, showing a cortical EEG electrode and a 32-channel silicon probe. **(E)** Examples of local field potentials (LFPs) and simultaneously recorded single unit activities (SUAs) during NREM (*left*) and REM sleep (*right*). Processed LFP signals for P-wave detection (red) are also shown. **(F)** Average waveforms of P-waves. *blue*, P-waves during NREM sleep; *green*, P-waves during REM sleep. Errors indicate SD. **(G)** Detected individual P-waves represented in first two PC (principal component) space. **(H)** Comparisons of average waveforms of P-waves (±50 ms from the trough of P-waves) between sleep states based on Pearson’s correlation coefficient (*n* = 20, *r* = 0.97 ± 0.01).

In all recording sessions where we could confirm REM sleep, we observed P-waves (**Figures 1C and E**). The position of electrodes was histologically confirmed in the mesopontine region (**Figure 1B**). P-waves often appeared as a cluster during REM sleep whereas P-waves during NREM sleep appeared less, typically in isolation (**Figure 1C**). P-waves during both sleep states were accompanied by spiking activity across simultaneously recorded brainstem neurons (**Figure 1E**), indicating that P-waves were locally generated neural events, rather than artifacts due to volume conduction or other electrical events. Thus, we could confirm P-waves in mice.

Since we also detected P-waves during NREM sleep, not just during REM sleep, we assessed whether detected P-waves were comparable between NREM sleep and REM sleep (**Figures 1F-H**). As shown in **Figure 1F**, the average waveform of P-waves was similar between states. To quantify this tendency, we applied principal component analysis (PCA) (**Figure 1G**) and measured Pearson’s correlation coefficient (**Figure 1H**). We confirmed the large overlap of individual waveforms between NREM and REM sleep in the first 2 PC space (**Figure 1G**) as well as the high correlation coefficient (0.975 ± 0.006) (**Figure 1H**), indicating that detected P-waves are quantitatively similar between sleep states.

### Similarity and difference in the basic properties of P-waves between sleep states

Next, we examined if the characteristics of P-waves are different between NREM and REM sleep (**Figure 2**). We began by comparing the frequency of P-waves between sleep states (**Figure 2A and Supplementary Figure 2**). The frequency of P-waves during REM sleep was around 0.5 – 1 Hz (0.81 ± 0.10 Hz), and this was significantly higher than NREM sleep (0.09 ± 0.01 Hz) (n = 38, *p* < 0.0001, signed-rank test). We did not find any statistically significant difference in P-wave frequency between recording methods (bipolar electrodes v.s. silicon probes; 31 v.s. 7) (*p* = 0.36 during NREM; *p* = 0.18 during REM, rank sum test with Bonferroni correction). Although our detection method could also detect P-wave-like events during wakefulness (**Supplementary Figure 2**), we focused on P-waves during NREM and REM sleep in this study.

**Figure 2.**
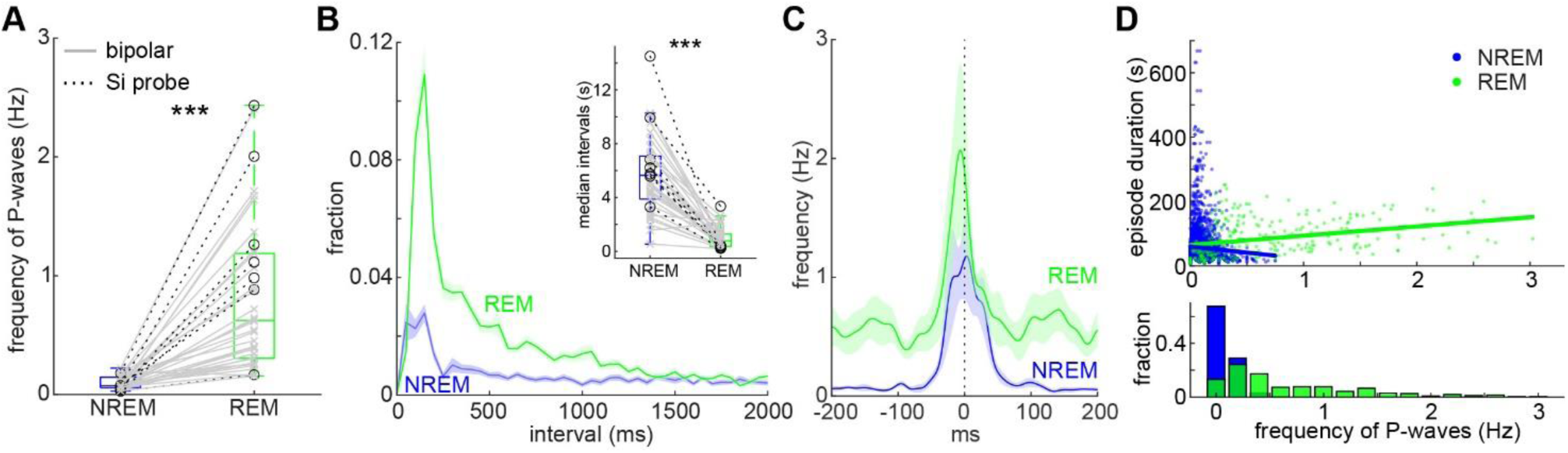
Features of P-waves during NREM and REM sleep. **(A)** Comparison of P-wave frequency between sleep states. *n* = 38 recordings. *** *p* <0.0001, signed-rank test. **(B)** Comparison of inter-P-wave intervals between sleep states. inset, comparison of the median intervals of P-waves between states. *n* = 38 recordings, *** *p* <0.0001, signed-rank test. **(C)** Cross-correlation of bilateral P-wave timing. *n* = 9 recordings. **(D)** The relationship between P-wave frequency and sleep episode duration. Lines are linear regression lines. *Bottom*, distributions of P-wave frequency during NREM (blue) and REM sleep (green).

Since P-waves during REM sleep often appear as clusters (**Figure 1**), we quantified the inter-event intervals (**Figure 2B**). As expected, a large fraction of P-waves appeared within 200 ms, especially during REM sleep. The median interval of P-waves was also significantly shorter during REM sleep (**Figure 2B *inset***) (n = 38, *p* < 0.0001, signed-rank test). To examine whether P-waves can occur synchronously in two hemispheres, we took advantage of some datasets (n = 9 sessions from 5 animals) where bipolar electrodes were implanted bilaterally (**Figure 2C**). Intriguingly, cross-correlation analysis showed the synchronous occurrence of P-waves between two hemispheres during both REM and NREM sleep. In addition, the cross-correlation profile during REM sleep exhibited a weak modulation in the 100 – 200 ms range, suggesting rhythmic theta synchrony between two hemispheres.

Because the frequency of P-waves gradually increases at around the onset of REM sleep as reported before ^15^, we wondered if the frequency of P-waves during REM sleep episode can correlate with the duration of REM sleep episodes (**Figure 2D**). We found that this is the case (*p* < 0.0001, *β* = 58.9, 95% confidence interval [45.1, 72.7], linear mixed-effects model) whereas the frequency of P-waves during NREM sleep did not correlate with episode duration (*p* = 0.87, *β* = -2.46, 95% confidence interval [-32.7, 27.8]). Altogether, although the waveform of P-waves is similar between sleep states, basic characteristics are markedly different between states.

### Similarity and difference in underlying brainstem neural activity between sleep states

While P-waves consist of a large deflection of LFPs (**Figure 1**), it remains unclear whether the underlying neural activity also consists of synchronous population firing or diverse neural activities. To address this issue, we analyzed single unit activity in the brainstem (**Figure 3**). Indeed, many neurons exhibited the peak of their activity around P-wave timing (**Figure 3A**). To examine whether the temporal structure of neural firing is preserved between NREM and REM sleep, we measured Pearson’s correlation coefficient between activity patterns in both states across neurons (**Figure 3B**). The mean of Pearson’s correlation coefficient was significantly higher than zero (*p* < 0.0001, *t*-test). Thus, the firing pattern associated with P-waves was generally preserved at the population level between sleep states.

**Figure 3.**
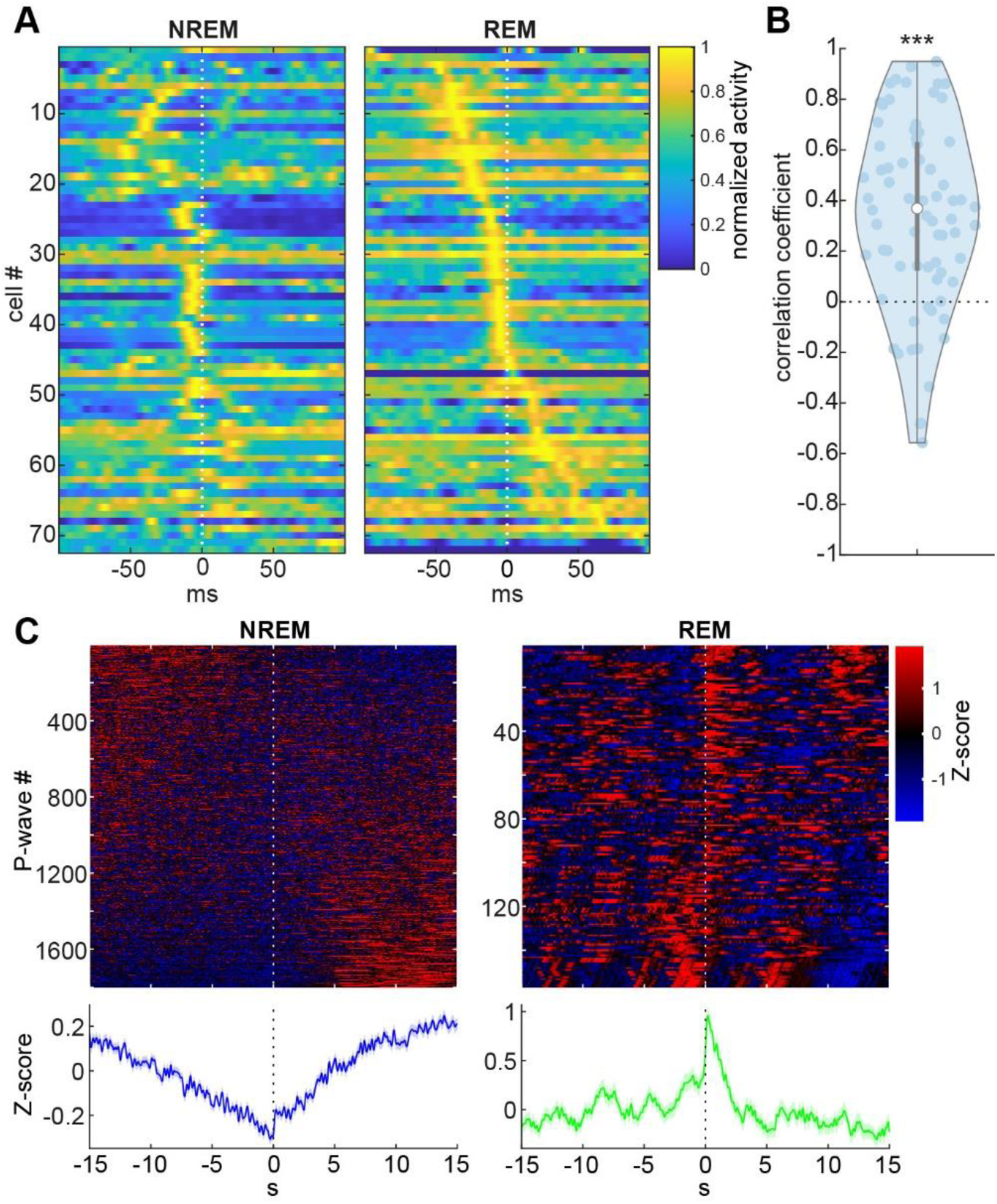
Brainstem neural firing associated with P-waves. **(A)** Normalized average firing profiles across cells during NREM (*left*) and REM sleep (*right*). Cells were sorted by peak timing during REM sleep. **(B)** Comparisons of mean firing profiles between sleep states across cells (n = 72) based on Pearson’s correlation coefficient. *** *p* < 0.0001, *t*-test. **(C)** Normalized (Z-scored) photometry signals of mesopontine cholinergic neurons associated with P-waves during NREM (*left*) and REM sleep (*right*). The top panels show individual normalized signal traces sorted by the 1^st^ PC score. The color range is between -1.96 and 1.96 (equivalent to 0.05 *p*-value). The bottom panels show the average of the normalized signals.

Since mesopontine cholinergic neurons have been implicated in P-wave genesis ^7, 8, 19^, we performed simultaneous electrophysiology and fiber photometry by expressing GCaMP6s in mesopontine cholinergic neurons. As we showed previously ^14^, P-waves were associated with the transient activity of cholinergic neurons during REM sleep (**Figure 3C and Supplementary Figure 3**). Although the transient activity of cholinergic activity was also confirmed during NREM sleep (**Supplementary Figure 3**), the level of cholinergic activity gradually decreased before P-waves and increased after P-waves (**Figure 3C**). We also confirmed similar state-dependent dynamics based on neural activity recorded with silicon probes (**Supplementary Figure 4**). Thus, although brainstem population activity associated with P-waves is generally similar between sleep states at a short (sub-second) timescale, at least mesopontine cholinergic neurons exhibit distinct activity patterns at a long (10s of seconds) timescale.

### Functional coupling between P-waves and hippocampal theta oscillations during REM sleep

Previously we demonstrated state-dependent functional coupling between P-waves and hippocampal oscillations across sleep states ^15^ and similar results were reported in macaque monkeys ^16^. However, it remains to be determined how hippocampal oscillations are modulated by coupling with P-waves across sleep states. In particular, it is unclear how sharp wave-ripples (SWRs) can be modulated by P-waves. To address these issues, we performed silicon or Neuropixels probe recording from the hippocampal CA1 while monitoring P-waves (**Figure 4A**). Although the Neuropixels probe allows monitoring of neural activity across multiple brain regions, in this study, we focus on the CA1 since the CA1 is the source of ripples ^3, 36, 37^.

**Figure 4.**
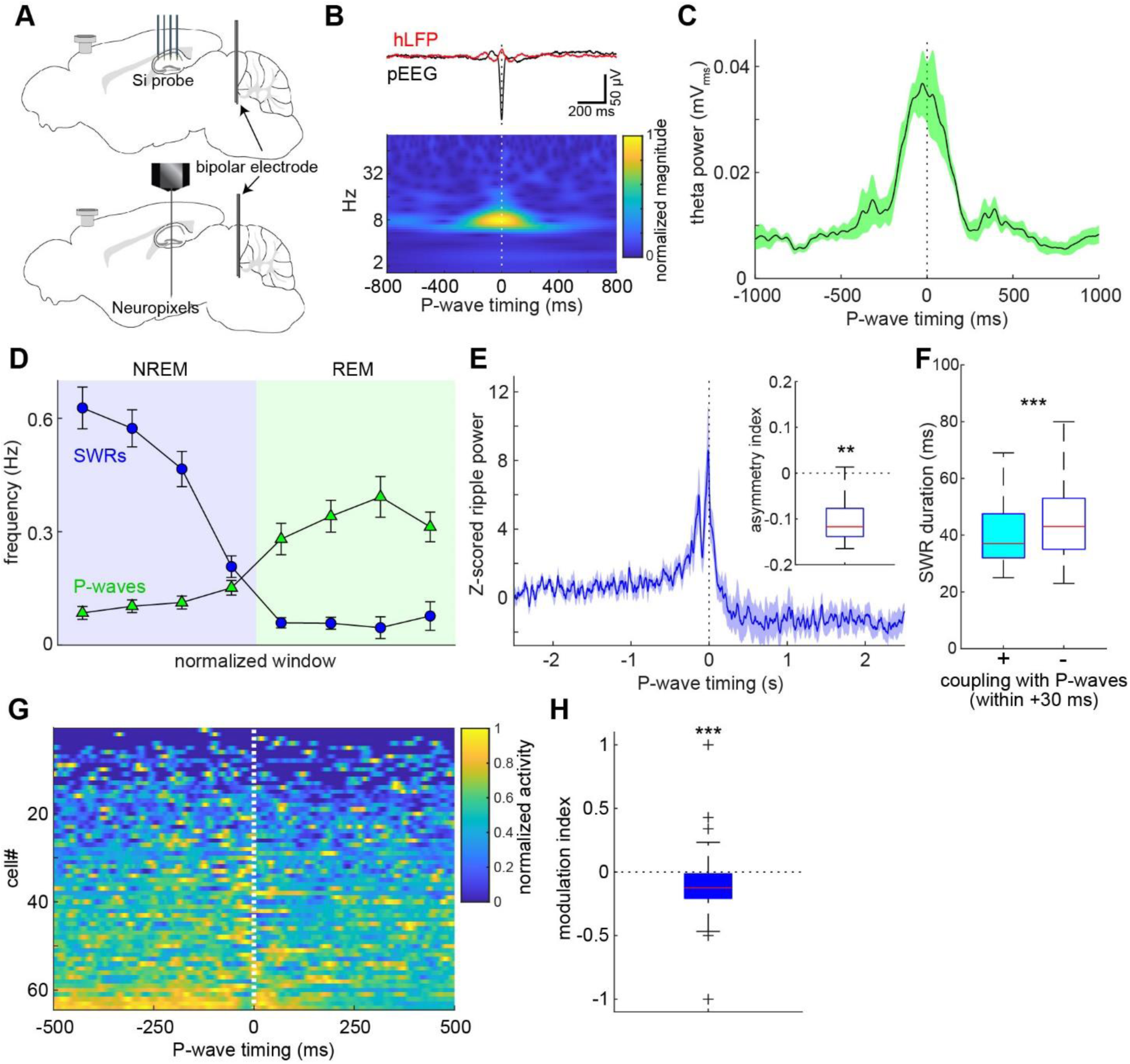
Functional coupling between P-waves and hippocampal oscillations during REM and NREM sleep. **(A)** A diagram of the experimental approach to hippocampal population recording. **(B)** *top*, the average hippocampal LFPs triggered by P-wave timing. *bottom*, a scalogram of P-wave-triggered hippocampal LFPs. **(C)** The average of hippocampal theta (4-10 Hz) power relative to P-wave timing. Event-triggered theta power was computed across recordings (n = 7). **(D)** Temporal profiles of SWRs (blue) and P-waves (green) across NREM and REM sleep episodes. Each episode duration was divided into four discrete time bins as normalized time windows. **(E)** Cross-correlation between the ripple power (80-250 Hz) profile and P-waves during NREM sleep. Each cross-correlation profile was z-scored based on the profile in a -3∼-2 sec window. *Inset*, the asymmetry index, a normalized difference in the normalized ripple power before and after P-wave timing in a 250 ms window. The negative value indicates the lower ripple power after P-waves. ** *p* < 0.005, *t*-test. n = 7. **(F)** Comparison of SWR duration between SWRs coupled with P-waves within 30 ms (n = 84) and other SWRs (n = 16689). *** *p* < 0.001, rank sum test. **(G)** Mean activity profiles of CA1 neurons (n = 65) relative to P-wave timing during NREM sleep. Cells were sorted by their first principal component score. **(H)** The modulation index, a normalized difference in firing rate before and after P-waves in a 250 ms window. *Plus signs*, outliers. *** *p* < 0.001, *t*-test.

Firstly, we confirmed the coupling between P-waves and hippocampal theta oscillations during REM sleep (**Figures 4B and C**). As expected, the average LFP signals showed rhythmic modulation around P-wave onset (**Figure 4B**). We assessed this in the frequency domain by applying a wavelet transformation (**Figure 4B**). This modulation was associated with an increase in their frequency band. We also examined when P-waves appeared during each theta cycle (**Supplementary Figure 5**). Significant phase coupling between theta oscillations and P-waves was observed across experiments (6 out of 7 recordings). Probably due to the gradual shift of theta phase across the CA1 region ^38^, the phase preference varied across experiments. On the other hand, we observed a consistent increase in theta power (**Figure 4C**). This increased theta power around P-wave timing was statistically confirmed by comparing the mean power between -1000 – -500 ms and -250 – 250 ms time windows (n = 7, *p* < 0.0001, *t*-test). These results reassure us that P-waves are coupled to hippocampal theta oscillations during REM sleep.

### Diminishing effects of P-waves on hippocampal SWRs during NREM sleep

Next, we examined if hippocampal SWRs coupled with P-waves have distinct features (**Figures 4D-F**). We began by comparing the frequency of SWRs and P-waves during NREM to REM sleep transitions (**Figure 4D**). These two neural events appeared complementary: while SWRs appeared mostly during NREM sleep (n = 7, 0.54 ± 0.04 Hz in NREM vs 0.04 ± 0.01 Hz in REM, *p* < 0.05, *t*-test), P-waves appeared more during REM sleep (n = 7, 0.09 ± 0.02 Hz in NREM vs 0.36 ± 0.09 Hz in REM, *p* < 0.05, *t*-test).

Although we previously reported that P-waves are coupled with SWRs ^15^, it is unclear if and how P-waves can be associated with the modulation of SWRs. To address this, we computed ripple power (80-250 Hz) around P-wave onset (**Figure 4E**). We found that ripple power sharply increases just before the timing of P-waves, consistent with our previous finding to demonstrate the coupling between SWRs and P-waves. However, ripple power rapidly decreased immediately after P-wave timing. This observation was confirmed across recordings (n = 7) by computing the asymmetry index that quantifies the difference in ripple power in a 250-ms window before and after P-waves (**Figure 4E *inset* and Supplementary Figure 6**). These results imply that the occurrence of P-waves may lead to the termination of SWRs.

To test this prediction, we compared the duration of SWRs depending on whether P-waves were coupled with SWRs in a short time window. While the fraction of SWRs coupled with P-waves within 30 ms was limited (0.6 ± 0.3 % of all SWRs during NREM sleep; 2.3 ± 0.9 % of all P-waves during NREM sleep, n = 7 recordings), we found that these SWRs coupled with P-waves were significantly shorter than the remaining SWRs (**Figure 4F**) (*p* < 0.0001, rank sum test). This tendency was confirmed within a 20 ─ 60 ms window (**Supplementary Figure 7**). We further examined whether CA1 neural activity is modulated by P-waves (**Figure 4G**). A group of neurons showed strong modulation at around the timing of P-waves: a subset showed a reduction in their firing after P-waves (**Figure 4G and Supplementary Figure 8**). To quantify this observation, we computed the normalized difference in firing rate before and after P-waves as the modulation index (**Figure 4H**). Overall CA1 neural firing was significantly reduced after P-waves during NREM sleep (*p* < 0.001, *t*-test). We further examined the directionality of coupling between hippocampal and pontine LFPs to confirm state-dependent functional coupling by calculating the spectral dependency ratio (**Supplementary Figure 9**). These results suggest that P-waves are functionally coupled to hippocampal oscillations in a sleep-state-dependent fashion. We uncovered the non-trivial relationship between P-waves and SWRs during NREM sleep.

## Discussion

Although P-waves have long been recognized as a signature of REM sleep, P-waves can also be observed during NREM sleep. However, the detailed characteristics and functional significance of P-waves during both sleep states remain poorly understood. Here, by performing multiple types of *in vivo* electrophysiological experiments and fiber photometry in mice, we described the similarity and difference in P-waves between sleep states in mice. Novel observations can be summarized as follows: first, P-waves appear synchronously in both hemispheres (**Figure 2C**). Second, while brainstem neural firing patterns associated with P-waves are generally similar between sleep states at a short timescale, mesopontine cholinergic activity shows distinct dynamics between sleep states at a longer timescale (**Figure 3**). Finally, we found that SWRs coupled with P-waves are short-lived (**Figure 4**).

### Comparisons with previous studies

P/PGO waves have been described in multiple mammalian species, including cats ^8–10, 12^, macaques ^16, 17^, rats ^13^ and humans ^18^. However, the existence of P-waves in mice has been anecdotal despite the importance of the animal model. While we recently showed P-waves in mice directly by using electrophysiological methods and indirectly by applying fiber photometry ^14, 15^, the characteristics of P-waves during NREM sleep were unclear.

P-waves in mice are similar to those in other species, with respect to the following features. First, the waveform of P-waves in mice (**Figure 1**) is similar to that of other species ^8^. In the present study, we quantitatively confirmed that the waveform of P-waves is comparable between REM and NREM sleep. Second, the temporal dynamics of P-waves in mice are also comparable with those in other species ^8^, with respect to the higher frequency of P-waves during REM sleep and a gradual increase in P-wave frequency around REM sleep onset.

In addition to these, we confirmed that P-waves are functionally coupled with SWRs and theta oscillations. A similar functional coupling between the hippocampus and cerebellum has been reported recently ^27^ although it remains to be confirmed if the neural events in the cerebellum are P-waves or not. More importantly, even macaque monkeys exhibit similar functional coupling ^16^. These results suggest that P/PGO waves are part of brain-wide neural ensembles across sleep states and sleep state-dependent functional coupling between P-waves and hippocampal oscillations is evolutionally preserved.

In the present study, we reported three non-trivial observations: first, P-waves appear synchronously in both hemispheres, implying an induction mechanism of P-waves (see below). Second, we showed that although P-waves are accompanied by mesopontine cholinergic activity during both sleep states, its dynamics at a long (over seconds) timescale is state-dependent: during NREM sleep, cholinergic activity gradually decreases before P-waves whereas cholinergic activity is at peak during P-wave genesis during REM sleep. Finally, although P-waves are functionally coupled with SWRs during NREM sleep, the occurrence of P-waves was associated with a reduction in ripple power as well as CA1 neural firing. These novel observations suggest the induction mechanism of P-waves as well as the functional significance of P-waves as discussed below.

### Possible mechanisms of P-wave genesis

The pons is known to be the origin of PGO/P-waves ^8, 39–41^. Our results are consistent with this notion: first, to detect P-waves, we subtracted LFP signals from adjacent channels to minimize the effect of volume conduction (**Figures 1C and E**). Second, we demonstrated that P-waves are accompanied by pontine spiking activity (**Figures 1E and 3A**).

In the present study, by performing extracellular electrophysiological recordings, we showed that brainstem neural ensembles consist of diverse functional classes while activity patterns are generally consistent between sleep states at a short (100s ms) timescale (**Figures 3A and B**), implying similar induction mechanisms of P-waves regardless of sleep states. In addition, the results from fiber photometry experiments (**Figure 3C and Supplementary Figure 3**) are consistent with the notion that mesopontine cholinergic neurons play a role in the induction of P/PGO waves ^8, 42^. Interestingly, we found that mesopontine cholinergic neurons exhibit distinct activity on a longer (>5 s) timescale between sleep states (**Figure 3C**), and a similar trend was also observed based on silicon probe recordings (**Supplementary Figure 4**), suggesting that pontine circuits involved in P-wave genesis may be influenced by distal activities in a state-dependent fashion.

In addition, P-waves appeared in both hemispheres synchronously (**Figure 2C**). Although we do not exclude the possibility that P-waves are generated in one hemisphere first, then P-waves start to synchronize in both hemispheres, our observation provides an insightful implication for the mechanism of P-wave genesis. Because a subset of mesopontine cholinergic neurons form commissural projections ^43^, these neurons may play a role in the synchronization of P-waves between two hemispheres. It would be worthwhile to investigate detailed anatomical and synaptic connections within the mesopontine region between hemispheres in the future.

### Functional implications of P-waves

What are the functional consequences of the state-dependent coupling of P-waves with hippocampal oscillations? Because both SWRs and theta oscillations have been implicated in systems memory consolidation ^3, 31, 44, 45^, P-waves may also play a role in memory consolidation. However, given our findings (**Figure 4**), the role of P-waves may also be state-dependent.

Although the exact role of REM sleep in systems memory consolidation remains controversial ^46, 47^, theta oscillations during REM sleep play a causal role in memory consolidation ^45^ and pharmacological manipulations of P-waves affect memory consolidation ^48^. While it has long been known that P-waves are phase-coupled with ongoing theta oscillations ^15, 16, 29, 30, 49, 50^, we also confirmed that P-waves can modulate hippocampal theta power. Because REM sleep can reorganize hippocampal population activity ^51^, it would be interesting to investigate how P-waves can contribute to such a reorganization of hippocampal excitability by manipulating P-waves.

Recently it has been shown that the duration of SWRs is causally linked to the degree of memory consolidation ^31^. Because we showed that SWRs coupled with P-waves are shorter than other SWRs, P-waves during NREM sleep may play a modulatory role to avoid excessive activation of hippocampal populations. A plausible mechanism of the shorter SWRs could be the influence of septal cholinergic neurons ^52, 53^. It remains to be determined how P-waves during NREM sleep can affect the downstream activity and how experience before sleep episodes can affect this unexpected, antagonistic interaction between SWRs and P-waves.

### Limitations of the study

Although our results provide insight into the functional significance of P-waves, the present study has limitations in the following aspects: first, determining the exact cell types of recorded neurons in the brainstem was impossible with silicon probe recording. Although we performed fiber photometry experiments to complement this limitation, this issue must be addressed by monitoring the activity of different cell types with genetically encoded indicators in the future. Second, related to the first point, although the electrode for P-wave monitoring was implanted in the medial parabrachial nucleus or nearby pontine regions, it may not have been optimal to monitor P-waves as we observed large variability in P-wave frequency across experiments (**Figure 2A**). It would be interesting to adopt a large-scale, high-density electrode array to systematically determine the origin of P-waves. Third, although we characterized state-dependent coupling between P-waves and hippocampal oscillations, it would be critical to directly determine the functional role of these couplings by manipulating P-waves while mice perform a behavioral task before and after sleep. Despite these limitations, however, our results provide a basis for future studies toward a better understanding of the regulatory mechanisms and functions of P-waves and sleep states.

### Conclusions

While P-waves appear synchronously in both hemispheres during both NREM and REM sleep, the frequency of P-waves differs between states. Underlying neural ensembles are generally consistent between states, suggesting a similar induction mechanism with state-specific, global activity patterns on a long timescale. Because the functional coupling between P-waves and hippocampal oscillations differs between sleep states, P-waves may contribute to brain-wide neural ensembles in a state-dependent fashion. In addition, state-dependent coupling between P-waves and hippocampal oscillations may lead to distinct consequences in their downstream neural circuits. Thus, these state-dependent coordinated activities may result in distinct functions of sleep states.

## Acknowledgements

We thank Dr Tomomi Sanagi, Dr Amisha Patel and Jacques Ferreira for technical assistance. This work was supported by Leverhulme Trust (RPG-2015-377), MRC (MR/V033964/1) and Horizon2020-ICT (DEEPER, 101016787) to SS, and by the Japan Society for the Promotion of Science (JSPS) Postdoctoral Fellow for Research Abroad, a Research Fellowship from the Uehara Memorial Foundation, FOREST (JPMJFR2047) from Japan Science and Technology Agency and JSPS KAKENHI (20H05047) to TT.

## Author contributions

TT and SM performed all *in vivo* electrophysiological experiments and associated histological analysis and sleep scoring. MM performed all Neuropixels probe experiments and pre-processing. SS performed all other data analysis. SS wrote the manuscript with inputs from TT, SM and MM.

## Disclosure Statement

The authors declare no competing financial interests. An earlier version of the manuscript has been posted on *bioRxiv* (https://doi.org/10.1101/2022.06.03.494781).

## Data Availability Statements

Quantitative data to reproduce figures are available along with low-level sleep scores and P-wave timing information online at https://doi.org/10.15129/d67d6b56-fa47-4f18-b244-002a74c504f8. All other data underlying this article will be shared on reasonable request to the corresponding author.

## Notes

### Competing Interest Statement

The authors have declared no competing interest.

### Summary of Updates

the title, Figures 1, 3 and 4, as well as main text revised.

