## Supplementary Data for "Pontine waves accompanied by short hippocampal sharp wave-ripples during non-rapid eye movement sleep"

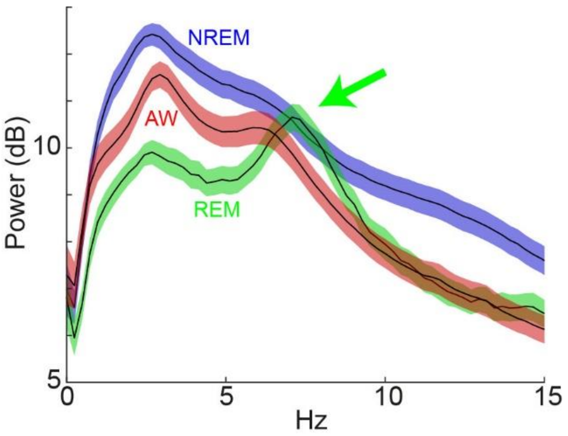


**Supplementary Figure 1. Average power spectral density in each state.**

High theta (~7 Hz) power was apparent during REM sleep (*arrow*). Note that due to the variability of low-cut filter settings across experiments, low frequency (<1 Hz) power did not reflect real signals. AW, wakefulness; NREM, NREM sleep; REM, REM sleep. Errors indicate SEM.

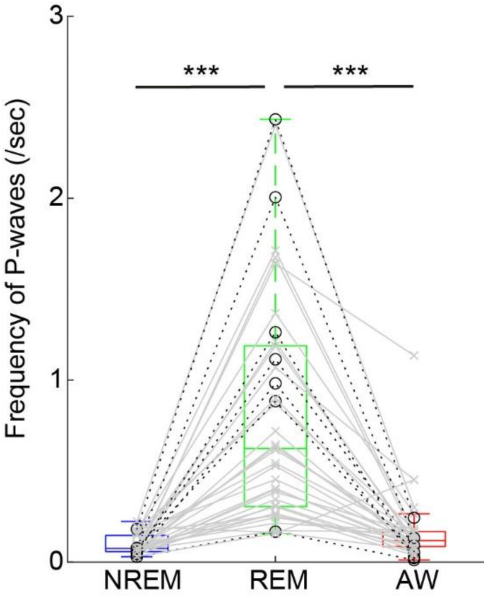


**Supplementary Figure 2. P-wave frequency across the sleep-wake cycle.**

*Circle*, silicon probe recording. *Cross mark*, bipolar electrode recording. ***, $p$ < 0.0001 (Kruskal-Wallis test with post-hoc Bonferroni method).

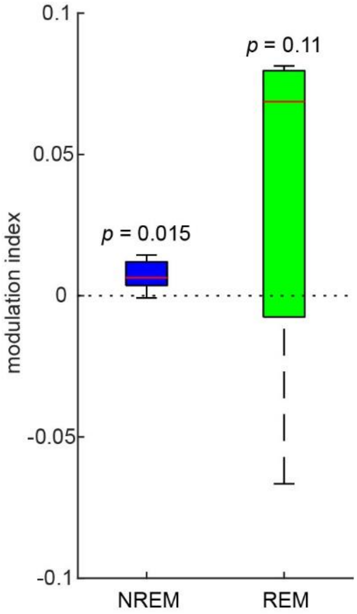


**Supplementary Figure 3**. **P-wave-associated calcium transients in mesopontine cholinergic neurons.**

The modulation index was defined as $(F_{post} - F_{pre})/(F_{post} + F_{pre})$, where $F_{pre}$ and $F_{post}$ are normalized fluorescent signals 1 s before and after P-wave timing, respectively. Wilcoxon signed-rank test was performed to examine if the median is zero.

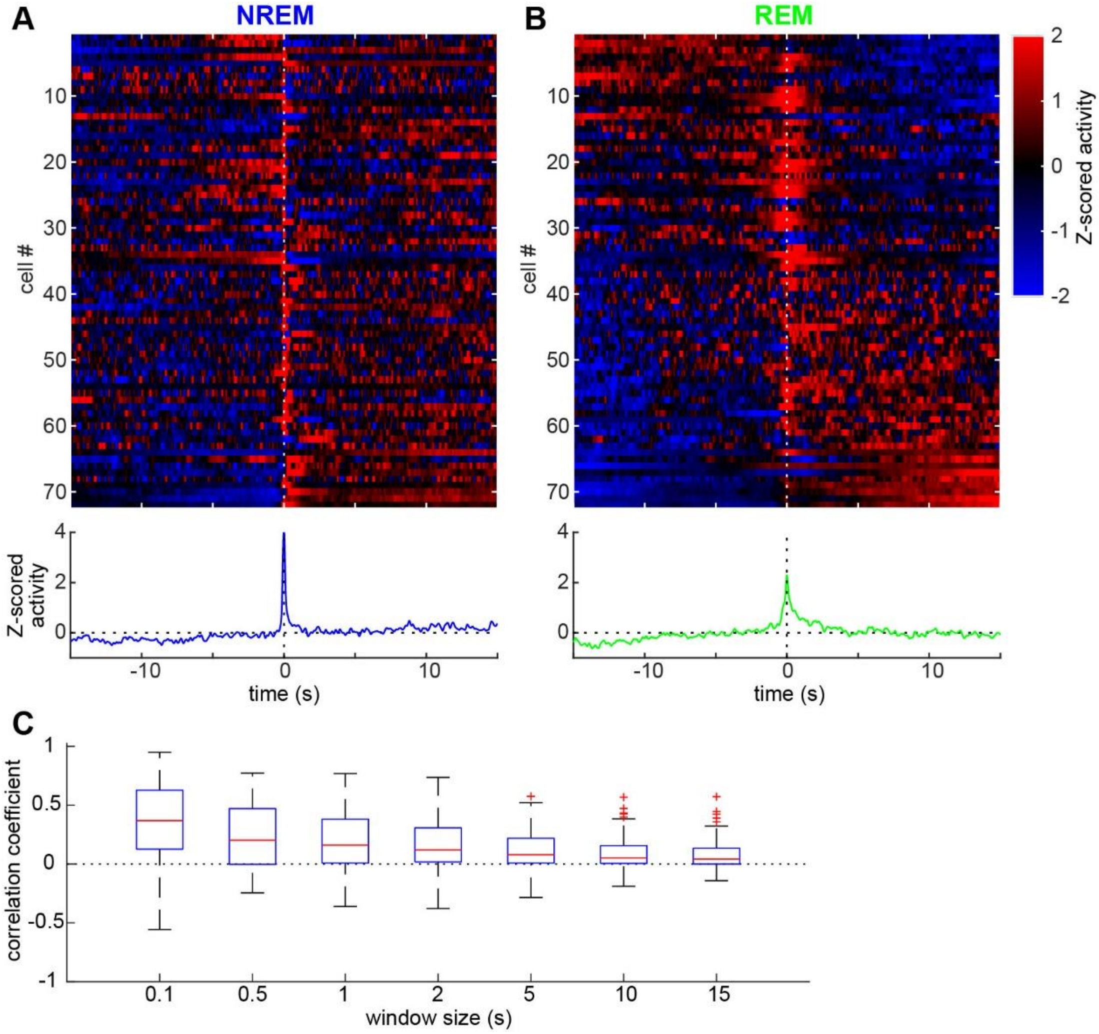


**Supplementary Figure 4**. **P-wave-associated brainstem neural ensemble dynamics on a long timescale.**

Brainstem neural activity during NREM sleep (**A**) and REM sleep (**B**). *top*, heat map of peri-event time histograms across recorded neurons. Neurons were sorted by the first principal component of the heat map during REM sleep. Individual neuronal activity was Z-scored. *bottom*, the average of Z-scored activity. (**C**) Pearson's correlation coefficient at different window sizes. As the window size increases, the correlation coefficient decreases ($p < 0.0001$, Kruskal-Wallis test).

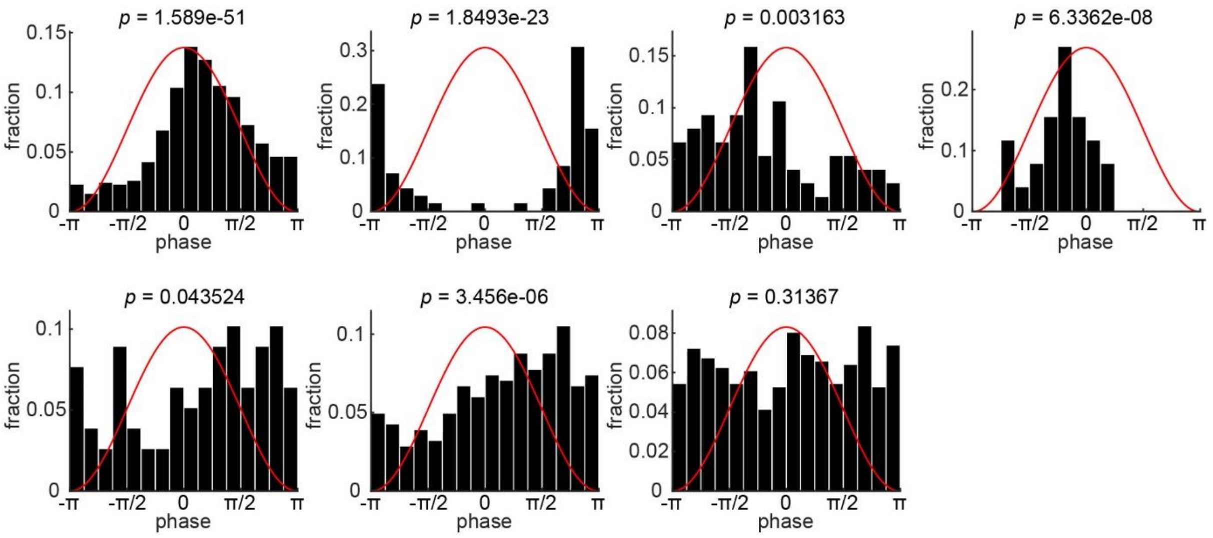


**Supplementary Figure 5. Theta-P-wave coupling during REM sleep.**

Phase histograms of P-wave timing relative to hippocampal theta rhythms across recordings (n = 7). *P*-values, Rayleigh's test. *Red line*, a theta cycle (4 – 10 Hz).

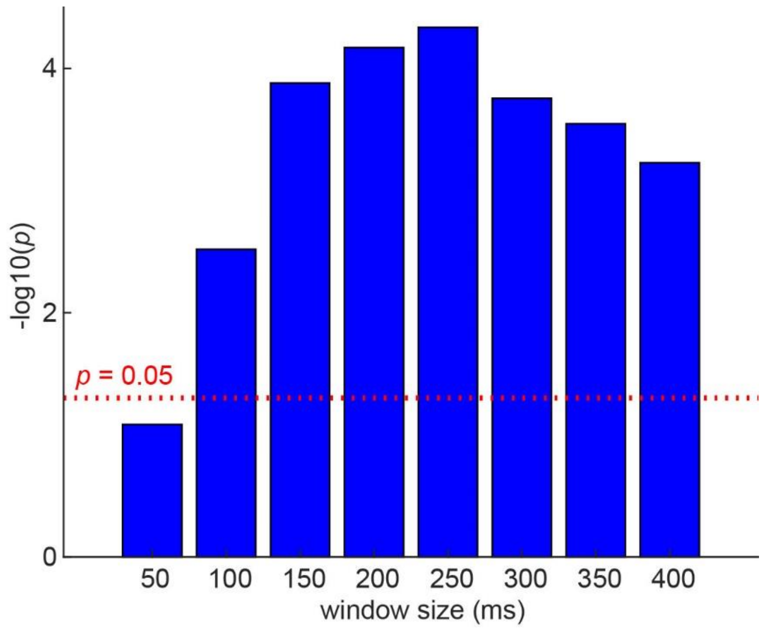


**Supplementary Figure 6. The effect of window size on ripple power change after P-waves.**

Related to **Figure 4E**, different window sizes were examined. After calculating the asymmetry index, *t*-test was performed to examine if the mean is zero. P-values were shown as a function of window size.

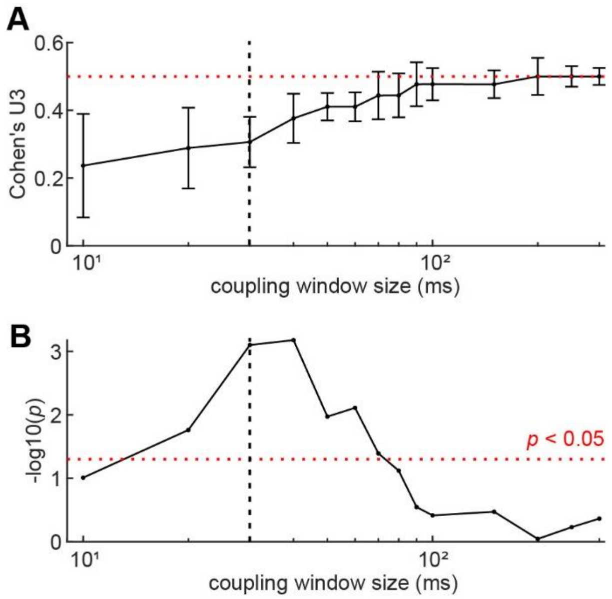


**Supplementary Figure 7. The relationship between coupling window size and statistical assessment.**

(**A**) Cohen's U3 as a function of coupling window size. Errors indicate 95% confidence intervals. (**B**) *P*-values (rank-sum test) as a function of coupling window size.

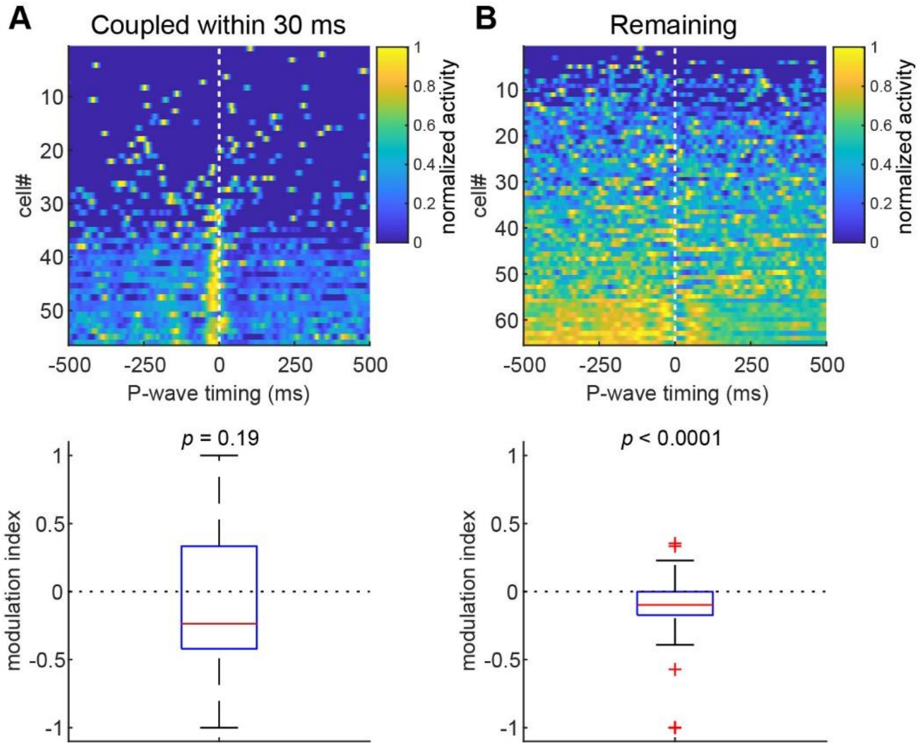


**Supplementary Figure 8. CA1 firing at P-wave timing depending on SWR coupling.**

CA1 firing at P-wave timing was separated depending on coupling to SWRs within a 30 ms (**A**) or longer time window (**B**). *top*, heat map of normalized peri-event time histograms across neurons. Individual neural activity was normalized by peak firing rate to 1. Neurons were sorted by the first principal component in each heat map. *bottom*, the modulation index was defined as $(FR_{post} - FR_{pre})/(FR_{post} + F_{Rpre})$, where $FR_{pre}$ and $FR_{post}$ are normalized firing rate 250 ms before and after P-wave timing, respectively. *t*-test was performed to examine if the mean is zero.

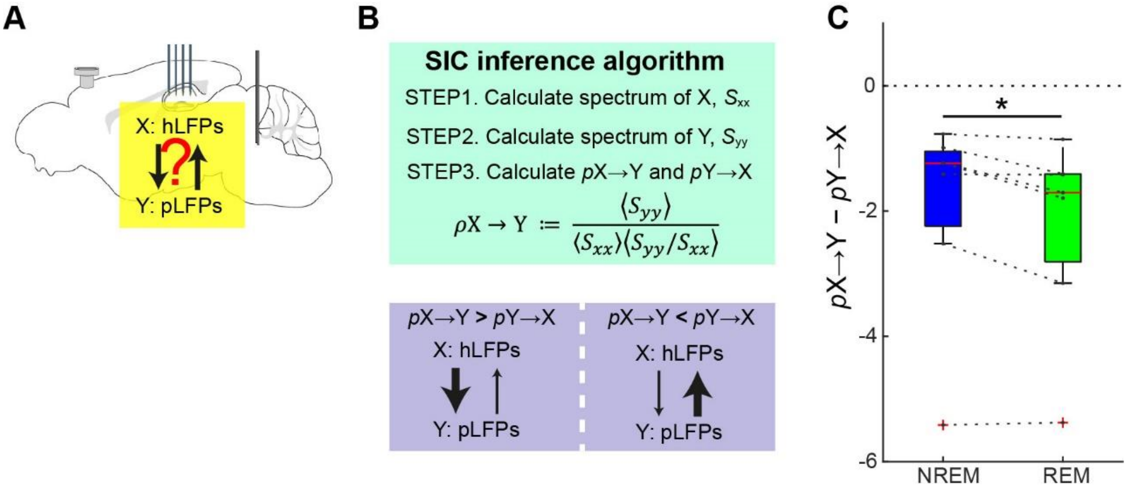


**Supplementary Figure 9. Spectral Independence Criterion analysis between hippocampal and pontine LFPs.**

(**A**) A schematic drawing outlines the motivation for this analysis. (**B**) An algorithm to compute the spectral density ratio based on the Spectral Independence Criterion (SIC). The interpretation of the outcome is also shown at the bottom. (**C**) The difference between *p*X→Y and *p*Y→X. The smaller value implies a stronger causal direction from pontine to hippocampal LFPs. *, $p < 0.05$ (signed rank test).
